# Prophylactic intranasal administration of a TLR2 agonist reduces upper respiratory tract viral shedding in a SARS-CoV-2 challenge ferret model

**DOI:** 10.1101/2020.09.25.309914

**Authors:** Pamela C. Proud, Daphne Tsitoura, Robert J. Watson, Brendon Y Chua, Marilyn J. Aram, Kevin R. Bewley, Breeze E. Cavell, Rebecca Cobb, Stuart Dowall, Susan A. Fotheringham, Catherine M. K. Ho, Vanessa Lucas, Didier Ngabo, Emma Rayner, Kathryn A. Ryan, Gillian S. Slack, Stephen Thomas, Nadina I. Wand, Paul Yeates, Christophe Demaison, David C. Jackson, Nathan W. Bartlett, Francesca Mercuri, Miles W. Carroll

**Affiliations:** National Infection Service, Public Health England (PHE), Porton Down, Salisbury, Wiltshire, United Kingdom SP4 0JG; Ena Respiratory, Level 9, 31 Queen St, Melbourne, Victoria, 3000, Australia; Department of Microbiology and Immunology, The University of Melbourne, at The Peter Doherty Institute for Infection and Immunity, 792 Elizabeth St, Melbourne, Victoria 3000, Australia; Viral Immunology and Respiratory Disease group and Priority Research Centre for Healthy Lungs, University of Newcastle and Hunter Medical Research Institute, Newcastle, Australia; Nuffield Dept of Medicine, Oxford University, Oxford, UK

**Keywords:** Ferret, COVID-19, SARS-CoV-2, animal model, coronavirus, viral shedding, TLR-2, transmission, INNA-051

## Abstract

Respiratory viruses such as coronaviruses represent major ongoing global threats, causing epidemics and pandemics with huge economic burden. Rapid spread of virus through populations poses an enormous challenge for outbreak control. Like all respiratory viruses, the most recent novel human coronavirus SARS-CoV-2, initiates infection in the upper respiratory tract (URT). Infected individuals are often asymptomatic, yet highly infectious and readily transmit virus. A therapy that restricts initial replication in the URT has the potential to prevent progression of severe lower respiratory tract disease as well as limiting person-to-person transmission.

We show that prophylactic intra-nasal administration of the TLR2/6 agonist INNA-051 in a SARS-CoV-2 ferret infection model effectively reduces levels of viral RNA in the nose and throat. The results of our study support clinical development of a therapy based on prophylactic TLR2/6 innate immune activation in the URT to reduce SARS-CoV-2 transmission and provide protection against COVID-19.

---

Coronaviruses (CoV) are pleomorphic, positive-sense, single-stranded RNA-enveloped viruses, members of the *Coronoviridae* family, that mainly infect wild animals and cause mild disease^1^. In addition, seven human CoVs that belong to either the Alphacoronavirus- or Betacoronavirus-genus have now been identified. Four of these human CoVs usually cause a mild, upper respiratory tract illness (common cold). Another three novel human CoVs have emerged in the past two decades through transmission to humans via an intermediate animal host^2^, and caused outbreaks of significant respiratory morbidity and mortality: in 2003, Severe Acute Respiratory Syndrome (SARS) CoV in China^3^, in 2012 Middle Eastern Respiratory Syndrome (MERS) CoV in Saudi Arabia^4^ and more recently in December 2019, SARS-CoV-2, a novel CoV type identified in lower respiratory tract of patients presenting viral pneumonia in Wuhan, China^5^. Unlike the highly pathogenic SARS or MERS CoVs, SARS-CoV-2 infections have spread rapidly around the globe, causing broad spectrum respiratory symptoms, from very mild to severe, life-threatening disease (COVID-19) mostly in at risk populations such as the elderly and those with comorbidities.

As with other respiratory CoVs, SARS-CoV-2 primarily spreads via the airborne route, with respiratory droplets expelled by infected individuals^6^. Virus can be transmitted from symptomatic, as well as pre- or asymptomatic individuals^7,8^, with asymptomatic individuals being able to shed virus, and therefore being capable to transmit the disease, for longer than those with symptoms^9^. As with other respiratory viruses such as influenza, recent evidence suggests that, the epithelium of the upper respiratory tract (URT) is the initial site of SARS-CoV-2 infection^10,11^. This is consistent with the abundant nasal epithelial cell expression of the SARS-CoV-2 receptor, angiotensin-converting enzyme 2 (ACE2) and its decreasing expression throughout the lower respiratory tract^11^.

A topical treatment of the URT that boosts anti-viral immunity and restricts viral replication is a promising method to promote viral clearance, reduce viral shedding and transmission. The TLRs are key microbe-recognition receptors with a crucial role in activation of host defence and protection from infections and therefore attractive drug targets against infectious diseases^12–14^. Synthetic agonists of the intracellular viral DNA/RNA-recognising TLR molecules, TLR3, TLR7/8 and TLR9, are capable of boosting protective innate immune responses against respiratory viruses. However, their success in the clinic has been limited, due to short-duration of benefit or induction of adverse effects, related to the release of pro-inflammatory cytokines and activation of the type-1 Interferon pathway^15–18^. TLRs expressed on the cell surface such as TLR2 offer an alternative approach. TLR2 dimerizes with TLR1 or TLR6 to recognize a broad variety of commensal and pathogenic microbial molecules and its activation is tightly regulated to maintain immune homeostasis^19^. A series of novel synthetic molecules, named the INNA compounds, have been developed with TLR2/6 agonist properties. Importantly, TLR2/6 agonists of the INNA compound series do not directly activate Type-1 interferons (unpublished data).

Airways administration of INNA compounds has been shown to protect from lethal influenza virus infection, prevent viral transmission and secondary bacterial superinfections in mouse disease models^20–22^. Intranasal (i.n.) treatment with INNA compounds also reduces viral load and lung inflammation in mouse models of rhinovirus infection (unpublished data). The demonstrated prophylactic benefit is associated with fast TLR2/6-mediated up-regulation of a series of innate immune response elements in airway epithelial cells, defined by early, rapid expression of NF-κB-regulated anti-microbial genes, including IFN-λ and chemokines, that precede immune cell recruitment and support prolonged antiviral defence, suppresses viral load and virus-induced pulmonary inflammation (unpublished data).

To determine whether TLR2/6 agonists are also active against SARS-CoV-2, we used prophylactic i.n. administration of the novel compound INNA-051, in a SARS-CoV-2 challenge ferret model^23^.

Ferret challenge models are commonly used to understand human respiratory virus-induced diseases and to evaluate the efficacy of related vaccines and drugs^24,25^. Use of ferrets is appropriate in the case of SARS-CoV-2 infection, as they express the virus entry ACE2 receptor in their airways^26–28^ and SARS-CoV-2 i.n. inoculation in ferrets results in virus replication in the URT and dose-dependent viral shedding^23,28^.

In the present study, ferrets received two i.n. administrations of INNA-051 or vehicle-control (PBS), at 4 days before and 1 day prior to i.n. challenge with 5.0 × 10^6^ plaque-forming units (PFU) of SARS-CoV-2 in 1ml volume (day 0). Four groups (6 animals/group) were used: Group 1 received two 1 ml doses (100μg/mL) INNA-051 (high dose); Group 2 received two 1 ml doses (20μg/mL) INNA-051 doses (low dose); Group 3 received one 1ml dose (20ug/mL) INNA-051 at −4 days and one 1 ml dose INNA-051 (100μg/mL) at −1 day (mixed dose); Group 4 received two doses of vehicle (PBS) alone. After inoculation with SARS-CoV-2, ferrets were monitored for 12 days. In life samples were taken at days 1, 3, 5, 7, 10 and 12, with scheduled culls at days 3 (n=6) and end of study days 12-14 (n=18) (Fig 1A).

**Figure 1.**
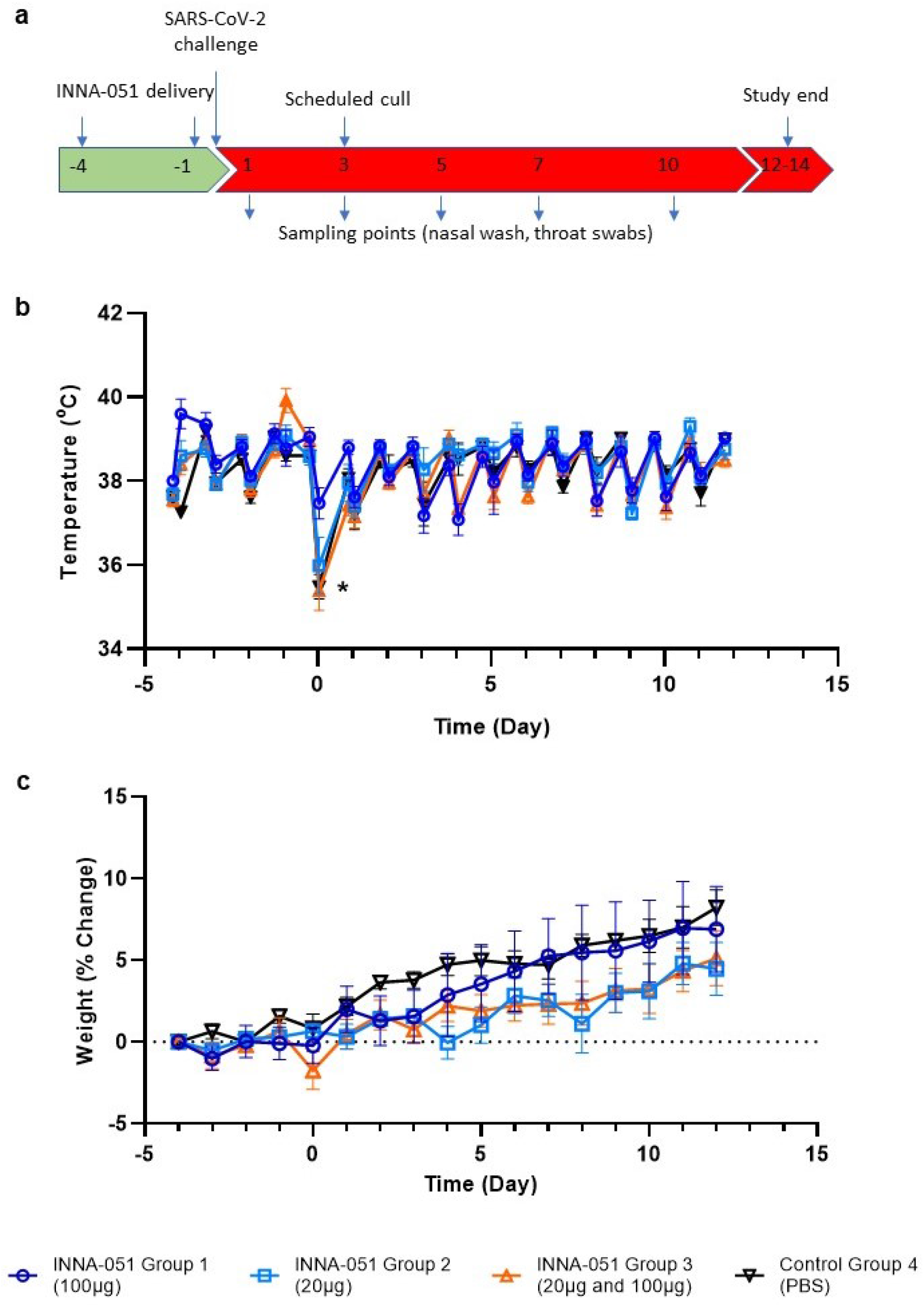
Clinical observations. (a) Schematic of experimental design. Ferrets received INNA-051 and PBS treatments 4 days and 1 day prior to challenge with 5.0 × 10^6^ pfu/ml SARS-CoV-2. Nasal wash and throat swabs were collected at days 1, 3, 5, 7, 10 & 12 post challenge (p.c.) for all treatment groups and control group. Scheduled culls were performed for 6/24 ferrets on day 3 p.c. and 18/24 ferrets on days 12-14 p.c. (b) Temperatures were measured twice daily (approximately 8 hours apart), using implanted temperature/ID chips. Mean temperatures +/− standard error of the mean (SEM) are displayed. Temperature dip post SARS-CoV-2 challenge (*) was attributed to sedation. (b) Weight was recorded daily and percentage change from the ferret weight prior to treatment plotted. Mean percentage weight change +/− SEM are displayed.

Previous *in vivo* studies in mice have shown that respiratory application of INNA compounds have a good safety profile, without significant pro-inflammatory side effects or systemic cytokine release syndrome (unpublished data). Intranasal administration to ferrets of two doses of INNA-051, prior to SARS-CoV-2 challenge, did not induce observable or, measurable clinical signs of inflammation or changes in the animal’s activity.

Assessment of body temperature revealed some variation between treatment groups (Fig 1B), with 2 of 6 ferrets in the INNA-051 high dose group 1 showing a transient increase of temperature >40.0°C, only after the first, but not the second dose. No significant changes from baseline were noticed in body weight, with the variation among groups being less than 2% (Fig 1C).

It has been previously described that SARS-CoV-2 infection in ferrets is not associated with the development of severe symptomatology, but it represents a robust model of mild disease that allows the study of respiratory viral replication^23,29^. In this context, no obvious differences in clinical disease signs were observed among treatment groups in this experimental study. Assessment of body temperature (Fig 1B) and weight loss (Fig 1C) did not reveal significant variation between the INNA-051 and PBS-treated groups.

To follow the dynamics of SARS-CoV-2 viral replication and assess the impact of INNA-051 prophylactic treatment, nasal wash and throat swab samples were taken 4 days before viral challenge and at 1, 3, 5, 7, 10 and 12-days post challenge (dpc). Analysis of viral RNA in nasal wash samples at 1 dpc confirmed infection in all treatment groups, with lower viral RNA levels detected in INNA-051 treatment Group 3 (Fig. 2A). Reduction of viral RNA in treatment Group 3 was also evident at 3 dpc (p=0.0155) (Fig. 2A). By 5 dpc. all INNA-051 treated groups had significantly (>10-fold) reduced viral RNA compared to the vehicle-control group (2-way ANOVA Dunnett’s multiple comparison test: Group 1 p=0.0244; Group 2 p=0.0107; Group 3 p=0.0071 compared to vehicle-control Group 4) (Fig 2A). On 5 dpc, the viral RNA levels in the nasal washes of the majority of INNA-051 treated animals remained low or below quantifiable limits throughout the course of infection. Viral RNA levels were found to be below the level of quantification in nasal washes of PBS-treated animals from 10 dpc onwards (Fig 2A).

**Figure 2.**
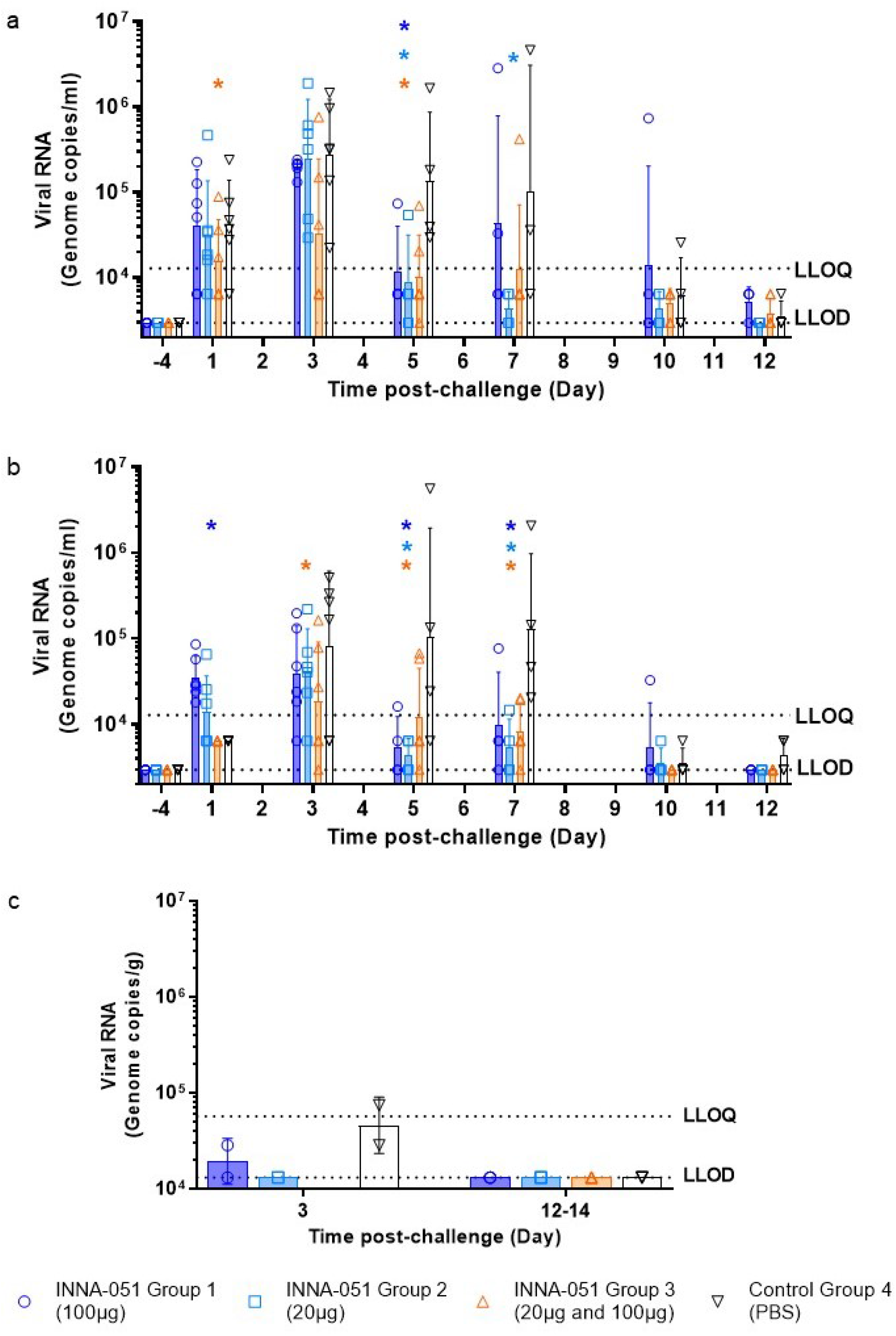
Viral RNA shedding following SARS-CoV-2 challenge. Nasal wash and throat swabs were collected at days 1, 3, 5, 7, 10 & 12 p.c. for all treatment groups and vehicle control group. Lung tissue was collected at necropsy on scheduled cull day 3 (n=6) and end cull days 12-14 (n=18). Viral RNA was quantified by RT-qPCR. (a) Nasal wash (b) Throat swab (c) Lung tissue. Geometric mean +/− standard deviation are displayed on the graphs. Dashed horizontal lines denote the lower limit of quantification (LLOQ) and lower limit of detection (LLOD). Statistical significance (95% CI of differences) in comparison to the control group using two-way ANOVA Dunnett’s multiple comparisons test are displayed above the error bars (*). Day 7 nasal wash for group 4 had viral RNA quantified for 3/4 ferrets; no sample was available for processing.

Analysis of viral RNA in throat swabs provided further evidence of the capacity of INNA-051 treatment to reduce SARS-CoV-2 in the URT (Fig 2B). On 3 dpc lower viral RNA levels were found in throat swabs of INNA-051 treated animals, with significantly greater reduction again observed (p=0.0345) in INNA-051 treatment Group 3. By 5 dpc, all groups treated with INNA-051 had significantly reduced (>10 fold) viral RNA levels, compared to the vehicle control group (2-way ANOVA Dunnett’s multiple comparison: Group 1 p=0.0002, Group 2 p=<0.0001 and Group 3 p=0.0039 compared to vehicle control Group 4). Highly significantly reduction in viral RNA in the throat of INNA-051 treated animals was also apparent on 7 dpc (2-way ANOVA Dunnett’s multiple comparison: Group 1 p=0.0014, Group 2 p=<0.0001 and Group 3 p=0.0002 compared to vehicle control Group 4), while by 10 dpc to the end of the study, the levels of viral RNA were below the limit of quantitation in all treatment groups (Fig 2B). Because all INNA-051 treatment groups exhibited reduced viral RNA in the nose and throat, we combined these groups into a single data set (supplementary figures) and compared to the group treated with vehicle. Using 2-way ANOVA Sidak’s multiple comparison test, significant (>10 fold) reduction in nasal viral RNA was observed at 5 dpc (p=0.0057) and highly significant (p<0.0001), greater than 10-fold reduction in throat viral RNA was apparent from 5 to 7 dpc following INNA-051 i.n. treatment (Figure S1). Group 2 (20ug/mL) appears to be the most optimal dosing in this study After 5 days post-exposure to SARS-CoV-2, INNA-051 had statistically significant reduction of virus in throat swabs (p=<0.0001) with 24 fold (96% reduction) and nasal wash (p=0.0107) 15 fold (93% reduction) in this group compared to untreated animals. These results indicate a similar profile with the protective effects of natural acquired immunity in the ferret challenge model, as observed following re-challenge with SARS-CoV-2^23^.

To assess SARS-CoV-2 detected beyond the URT, lung tissue samples were collected, on scheduled cull day 3 (6/24 animals) and days 12-14 (18/24 animals) dpc and analysed for viral RNA levels. On day 3 dpc, two culled ferrets from the control vehicle group had detectable viral RNA levels (7.42×10^4^ and 2.86×10^4^ copies/ml) (Fig 2C). There was one ferret in Group 1 showing detectable, but below the quantifiable limit, viral RNA, and no other INNA-051 treated ferrets showing detectable viral RNA in lung tissue on day 3 and days 12-14 dpc

This study provides evidence supporting a novel approach to prevent SARS-CoV-2 transmission, based on reduced viral shedding, following prophylactic i.n administration of INNA-051. Global efforts for prevention of SARS-CoV-2 infection have so far been mostly focused on social distancing and hygiene measures as well as on R&D efforts for the development of vaccines. Our data demonstrate, for the first time, in an *in vivo* SARS-CoV-2 infection model, that INNA-051 is highly effective at reducing URT viral shedding, providing the potential to control virus transmission and COVID-19 disease.

TLR2 stimulation at mucosal surfaces triggers rapid up-regulation of protective, innate immune defence responses, and also activates counter-regulatory signalling that suppresses development of excessive inflammation and tissue damage and promotes the integrity of local epithelial barrier function^30,31^. In addition, the INNA compounds have been specifically designed to exert TLR2-mediated pharmacological activity on mucosal epithelium, without being systemically absorbed (Ena Respiratory unpublished data), a property that is expected to facilitate their development as safe, antiviral drug candidates. The lack of obvious clinical signs of inflammation following the administration of two doses of INNA-051 administered i.n supports this view. Histopathology from the study indicates that i.n. INNA-051 administration does not exacerbate SARS-CoV-2 pathology in the ferret lung in this setting (Figure S2).

It has been previously shown that i.n administration of an INNA compound in a mouse model of influenza triggers a cascade of innate immune signals that results in reduction of viral load, prevention of lower-respiratory infection and viral transmission between animals^20–22^. In a mouse rhinovirus infection model, we have shown that prophylactic treatment with an INNA compounds primes airway mucosal immunity providing prolong protection (unpublished data). Effective and accelerated anti-viral responses against rhinovirus have also been demonstrated following INNA compound treatment in an *in vitro* model of primary bronchial epithelial cells from healthy individuals and donors with asthma (unpublished data).

Although the ferret SARS-CoV-2 model has limitations and may not represent the severe spectrum of COVID-19 disease, our findings are highly encouraging and indicative of the potential impact i.n. administration of INNA-051 prophylactically may have against SARS-CoV-2 in humans. The fact that a significant reduction of URT viral RNA levels was observed in INNA-051-treated outbred ferrets during the peak of viral replication (5-7 days dpc) in this model^23^ implies airway immunity priming and enhancement of antiviral host defence. The predictive value of antiviral effectiveness data from respiratory viral infection ferret models and translation into human infectious disease has been established^25,32^. For this reason, the SARS-CoV-2 ferret model has been used, during the current pandemic, to evaluate the therapeutic effect of a number of FDA-approved/repurposed drugs including, lopinavir-ritonavir, hydroxychloroquine sulfate, or emtricitabine-tenofovir^33^. These drugs were found to have no or only modest (~4 fold for emtricitabine-tenofovir) effect against SARS-CoV2 viral replication, as measured by viral titres in nasal wash from the ferrets^33^.

Substantial reduction of SARS-CoV viral shedding in the URT and therefore control of respiratory virus transmission may not be easily achievable without potentiation of airways antiviral immune defences^34^. Systemic antiviral drugs, as well as vaccines may not be effective in halting respiratory viral transmission even if they achieve suppression of clinical disease and in fact preliminary results from an experimental study with one of the leading SARS-CoV-2 vaccine candidates (an adenovirus-construct expressing SARS-CoV-2 spike protein) in non-human primates have shown little effect on the virus load in nasal washes^35^. To address these potential limitations, particularly during the urgent circumstances of an epidemic, parallel use of an i.n. administered innate immune modulator with the characteristics of INNA-051 may be highly appropriate to rapidly boost innate immunity at the primary site of respiratory infection which is protective within days of treatment. The use of i.n. INNA-051 for antiviral respiratory prophylaxis therefore offers several additional advantages, including fast-acting protection, and is in contrast to vaccines that take 2-4 weeks to mount a protective response. The limited risk for development of antiviral resistance, the option of self-administration and the non-prohibitive cost for large-scale manufacturing are also especially attractive factors.

In conclusion, this study provides evidence that prophylactic i.n. administration of the TLR2/6 agonist INNA-051 offers a promising approach for prevention and management of SARS-CoV-2 infection that can be used as a stand-alone method of antiviral prophylaxis and is complimentary to potential vaccination programs. This approach is particularly appealing to individuals at elevated risk of community transmission or development of severe disease, including front-line health care workers, vulnerable communities, the elderly, the immunocompromised and those with existing comorbidities.

## Contributions

N.W.B, B.Y.C, C.D, D.C.J, F.M, P.P, D.T and M.W.C. contributed to the concept and design of the study.

P.P and D.T. wrote the manuscript.

N.W.B, B.Y.C, C.D, S.D, D.C.J, F.M. and M.W.C. provided critical review.

P.P co-ordinated the study and performed analysis on data generated.

J.D. and M.G.C. provided virus strain.

K.R.B. grew viral stock, provided challenge material and assisted with statistical analysis.

S.A.F and P.Y. managed all animal procedures.

B.E.C, R.C, C.M.K.H, V.L, D.N, K.A.R. and S.T. contributed to the processing of all animal samples at containment level 3.

E.R directed histology experiments and performed critical assessment of pathology.

M.J.A, N.I.W and R.J.W. performed the extraction and PCR of samples.

G.S performed quality control and analytical assistance on PCR data.

## Acknowledgements

The authors gratefully acknowledge the support from the Biological Investigations Group at the National Infection Service, PHE, Porton Down, United Kingdom. Special thanks to Thomas Hender, Carrie Turner, Stephen Findlay-Wilson and Neil McLeod for assisting in providing RT-qPCR data for this work. The authors would like to express their gratitude to Jade Gouriet, Phillip Brown, Karen Gooch and Jemma Paterson for their help in processing of Containment Level 3 *in vivo* samples. The authors would like to thank Laura Hunter, Chelsea L. Kennard and Francisco J. Salguero for their contribution to the processing and critical review of pathology tissues. The authors are grateful to Michael G. Catton and Julian Druce from Victorian Infectious Disease Reference Laboratory for the generous donation of the SARS-CoV-2 strain.

## Methods

### Animals

Twenty-four healthy, female outbred ferrets (*Mustela putorius furo*) aged 6-8 months were obtained from a UK Home Office accredited supplier. The mean weight at the time of first INNA-051 treatment was 845g/ferret (range 740-1040g). Animals were housed in social groups of six prior to and post INNA-051 treatment at Advisory Committee on Dangerous Pathogens (ACDP) containment level 2. Animals were transferred to ACDP containment level 3 and housed in pairs post SARS-CoV-2 challenge. Cages met with the UK Home Office *Code of Practice for the Housing and Care of Animals Bred, Supplied or Used for Scientific Procedures* (December 2014). Access to food and water was *ad libitum* and environmental enrichment was provided. All experimental work was conducted in accordance with and under the authority of a UK Home Office approved project licence that had been subject to local ethical review at PHE Porton Down by the Animal Welfare and Ethical Review Body (AWERB) as required by the *Home Office Animals (Scientific Procedures) Act 1986*. Animals were sedated by intramuscular injection of ketamine/xylazine (17.9 mg/kg and 3.6 mg/kg bodyweight) for administering of treatments, in-life sampling and viral challenge.

### INNA-051

INNA-051 belongs to a series of closely-related, pegylated synthetic analogues of the diacylated lipopeptide, S-[2,3-bis(palmitoyl oxy)propyl] cysteine (Pam_2_Cys) (INNA compound series), with selective TLR2/TLR6 agonist activity. Pam_2_Cys is inherently insoluble and has been rendered soluble by others through addition of the amino acid motif SK4^36^. Oligo lysine sequences have, however, been shown to be toxic, albeit at high concentration^37^ and to modulate viral infection processes independent of TLR activation^38^. Any off-target effects were mitigated by incorporating polyethylene glycol as a solubilising agent, in the INNA compound series^38^. The EC50s for INNA-051 for the human TLR2/6 receptor is calculated at 40.1 pg/mL or ~19pM.

Freeze dried INNA-051 provided by Ena Respiratory, Melbourne, Australia was resuspended in phosphate buffered saline (PBS) (1mg/ml) and stored 2-8°C. Immediately prior to treatment, INNA-051 (1mg/ml) was further diluted in PBS to the required treatment doses; high dose (100μg/ml), low dose (20μg/ml) and mixed dose (20μg/ml first dose and 100μg/ml second dose).

### Inoculum

SARS-CoV-2 Victoria/01/2020^39^ was generously provided by Peter Doherty Institute for Infection and Immunity, Melbourne, Australia at P1 and passaged twice in Vero/hSLAM cells [ECACC 04091501], obtained from the European Collection of Authenticated Cell Cultures (ECACC) PHE, Porton Down, UK. Whole genome sequencing was performed, on the challenge isolate, SISPA protocol and then sequenced using Nanopore as described previously^40^. Virus titre was determined by plaque assay on Vero/E6 cells [ECACC 85020206]. Challenge substance dilutions were conducted in Phosphate Buffer Saline (PBS). Challenge virus was delivered by intranasal instillation (1.0 ml total, 0.5 ml per nostril) at 5.0 × 10^6^ pfu/ml.

### Experimental Design

Experimental design and viral challenge dose were informed by a previous dose-dependent ferret study^23^. Prior to commencing the experiment, animals were randomly assigned to the four treatment groups, to minimise bias. A temperature/ID chip (Bio-Thermo Identichip, Animalcare Ltd, UK) was inserted subcutaneously into the dorsal cervical region of each animal.

INNA-051 was delivered by intranasal instillation (1.0 ml total, 0.5 ml per nostril) to three groups (n=6) of ferrets 4 days and 1 day prior to challenge. On each day, group 1 received a high dose [100μg/ml], group 2 a low dose [20μg/ml] and group 3 received a 20μg/ml dose 4 days prior to challenge and a 100μg/ml] dose 1 day before challenge. PBS was delivered to control group ferrets (n=6) 4 days and 1 day prior to challenge. Two ferrets each from the high dose, low dose and control groups were scheduled for euthanasia on day 3 (n=6). Remaining ferrets (n=18) were scheduled for euthanasia on days 12-14; high and low dose [day 12 n=1, day 13 n=2, day 14 n=1], mixed dose [n=2 days 12-14] and control [n=2 days 12 and 14].

Nasal washes and throat swabs for all ferrets were taken prior to first treatment, at days 1 and 3 pc (n=24) and at days 5, 7, 10 and 12 pc for surviving ferrets (n=18). At necropsy, tissue samples were taken for histopathology and analysed by PCR. Nasal washes were obtained by flushing the nasal cavity with 2 ml PBS. Cotton throat swabs (Koehler Technische Produkten, VWR) were gently stroked across the back of the pharynx in the tonsillar area and retained in viral transport media (VTM). Throat swabs were processed, and aliquots were stored in AVL at −80°C until assay.

### Clinical and euthanasia observations

Animals were monitored for clinical signs of disease twice daily (approximately 8 hours apart) for the entirety of the experiment. Clinical signs of disease were assigned a score based upon the following criteria. Activity was scored as follows; 0 = alert and playful, 1 = alert, playful when stimulated, 2 = alert, not playful when stimulated, 3 = not alert or playful. No clinical signs were noted throughout the experiment. To meet the requirement of the project license, immobility, neurological signs or a sudden drop in temperature were predetermined automatic euthanasia criteria. Animals were also deemed to have reached a humane endpoint if their body weight was at or below 30% baseline. If any ferret reached any of these three criteria, they were to be immediately euthanised using a UK Home Office approved Schedule 1 procedure. No animals reached these end-points during this study.

Temperature was taken using a microchip reader and implanted temperature/ID chip. Temperature was recorded at each clinical scoring point using the chip to ensure any peak of fever was recorded. Animals were weighed at the same time each day throughout the experiment.

### Necropsy Procedures

Ferrets were anaesthetised with ketamine/xylazine (17.9 mg/kg and 3.6 mg/kg bodyweight) and exsanguination was effected via cardiac puncture, followed by injection of an anaesthetic overdose (sodium pentabarbitone Dolelethal, Vetquinol UK Ltd, 140 mg/kg). A necropsy was performed immediately after confirmation of death. The left lung was dissected and used for subsequent virology procedures.

### RNA Extraction

RNA was isolated from nasal wash, throat swabs, EDTA treated whole blood, BAL and lung tissue. Weighed lung tissue was homogenised and inactivated in RLT (Qiagen) supplemented with 1%(v/v) Beta-mercaptoethanol. Tissue homogenate was then centrifuged through a QIAshredder homogenizer (Qiagen) and supplemented with ethanol as per manufacturer’s instructions. Non-tissue samples were inactivated in AVL (Qiagen) and ethanol. Downstream extraction on all inactivated samples was then performed using the BioSprint™96 One-For-All vet kit (Indical) and Kingfisher Flex platform as per manufacturer’s instructions.

### Quantification of Viral Loads by RT-qPCR

Reverse transcription-quantitative polymerase chain reaction (RT-qPCR) targeting a region of the SARS-CoV-2 nucleocapsid (N) gene was used to determine viral loads and was performed using TaqPath™ 1-Step RT-qPCR Master Mix, CG (Applied Biosystems™), 2019-nCoV CDC RUO Kit (Integrated DNA Technologies) and QuantStudio™ 7 Flex Real-Time PCR System. Sequences of the N1 primers and probe were: 2019-nCoV_N1-forward, 5’ GACCCCAAAATCAGCGAAAT 3’; 2019-nCoV_N1-reverse, 5’ TCTGGTTACTGCCAGTTGAATCTG 3’; 2019-nCoV_N1-probe, 5’ FAM-ACCCCGCATTACGTTTGGTGGACC-BHQ1 3’. The cycling conditions were: 25°C for 2 minutes, 50°C for 15 minutes, 95°C for 2 minutes, followed by 45 cycles of 95°C for 3 seconds, 55°C for 30 seconds. The quantification standard was *in vitro* transcribed RNA of the SARS-CoV-2 N ORF (accession number NC_045512.2) with quantification between 1 × 10^1^ and 1 × 10^6^ copies/μl. Positive samples detected below the lower limit of quantification (LLOQ) were assigned the value of 5 copies/μl, whilst undetected samples were assigned the value of < 2.3 copies/μl, equivalent to the assays lower limit of detection (LLOD).

### Histopathological Analysis

Samples from the left cranial and left caudal lung lobe together with nasal cavity, were fixed by immersion in 10% neutral-buffered formalin and processed routinely into paraffin wax. Nasal cavity samples were decalcified using an EDTA-based solution prior to embedding. 4 μm sections were cut and stained with haematoxylin and eosin (H&E) and examined microscopically. In addition, samples were stained using the RNAscope technique to identify the SARS-CoV-2 virus RNA. Briefly, tissues were pre-treated with hydrogen peroxide for 10 minutes (room temperature), target retrieval for 15 mins (98-101°C) and protease plus for 30 mins (40°C) (Advanced Cell Diagnostics). A V-nCoV2019-S probe (Cat No. 848561, Advanced Cell Diagnostics) was incubated on the tissues for 2 hours at 40°C. Amplification of the signal was carried out following the RNAscope protocol using the RNAscope 2.5 HD Detection kit – Red (Advanced Cell Diagnostics).

### Statistical analysis

Virology data were compared by two-way ANOVA with Dunnett’s multiple comparisons test and two-way ANOVA with Sidak’s multiple comparisons test. Transformed values fitted to a straight line in a QQ plot confirming normal distributions for comparison. A p-value <0.05 was considered statistically significant. Statistical analyses were performed using GraphPad Prism, version 8.4.2.

