## Supplementary Figures for "Prophylactic intranasal administration of a TLR2 agonist reduces upper respiratory tract viral shedding in a SARS-CoV-2 challenge ferret model"

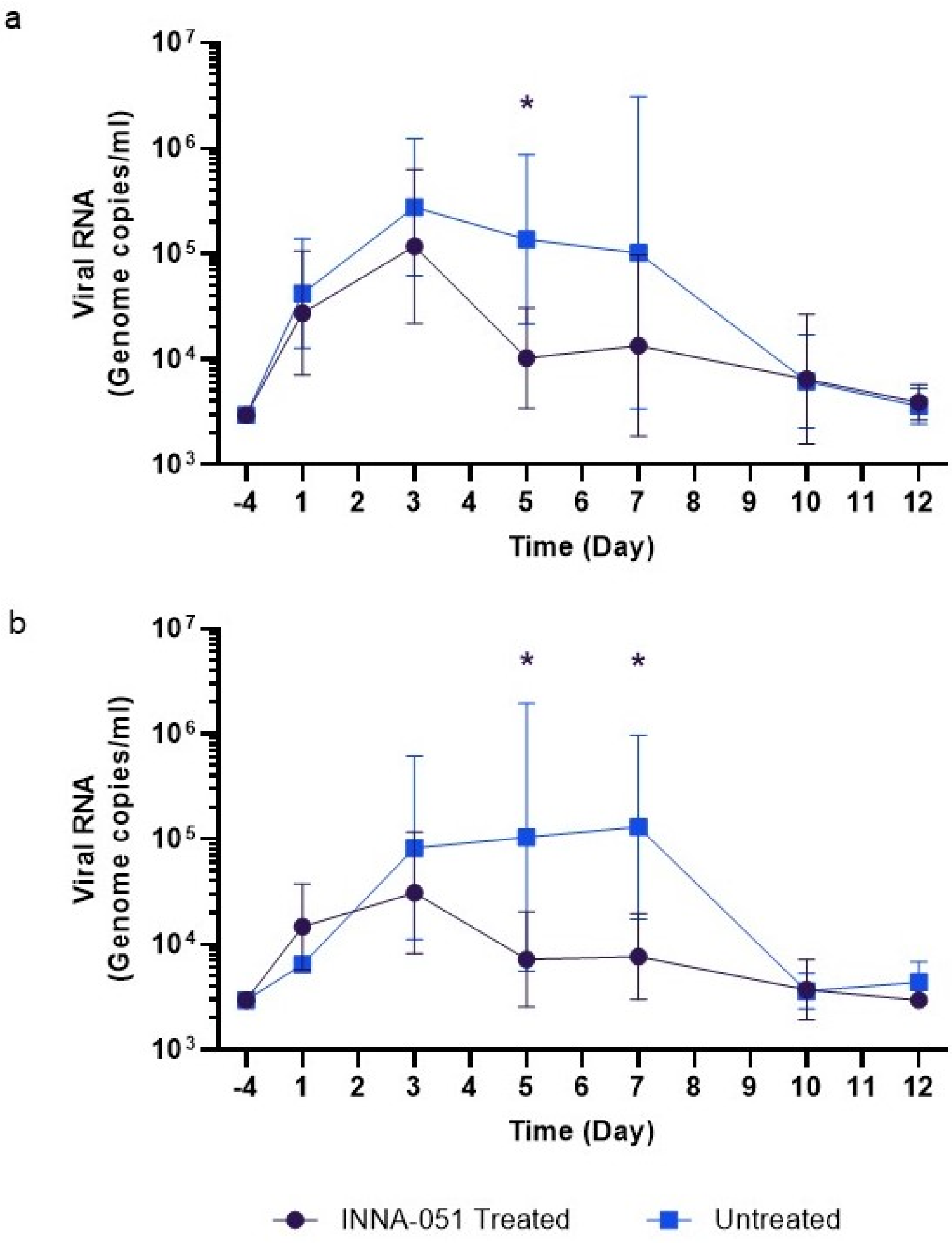


**Supplementary Figure 1. Viral RNA shedding following SARS-CoV-2 challenge.** Quantified viral RNA by RT-qPCR were combined for INNA-051 treated groups 1-3 and compared against untreated vehicle control group 4. (a) Nasal wash (b) Throat swab. Geometric mean +/- standard deviation are displayed on the graphs. Dashed horizontal lines denote the lower limit of quantification (LLOQ) and lower limit of detection (LLOD). Statistical significance (95% CI of differences) between groups using two-way ANOVA Sidak's multiple comparisons test are displayed above the error bars (*); nasal wash day 5 p=0.0057, throat swab days 5 and 7 p=<0.0001.

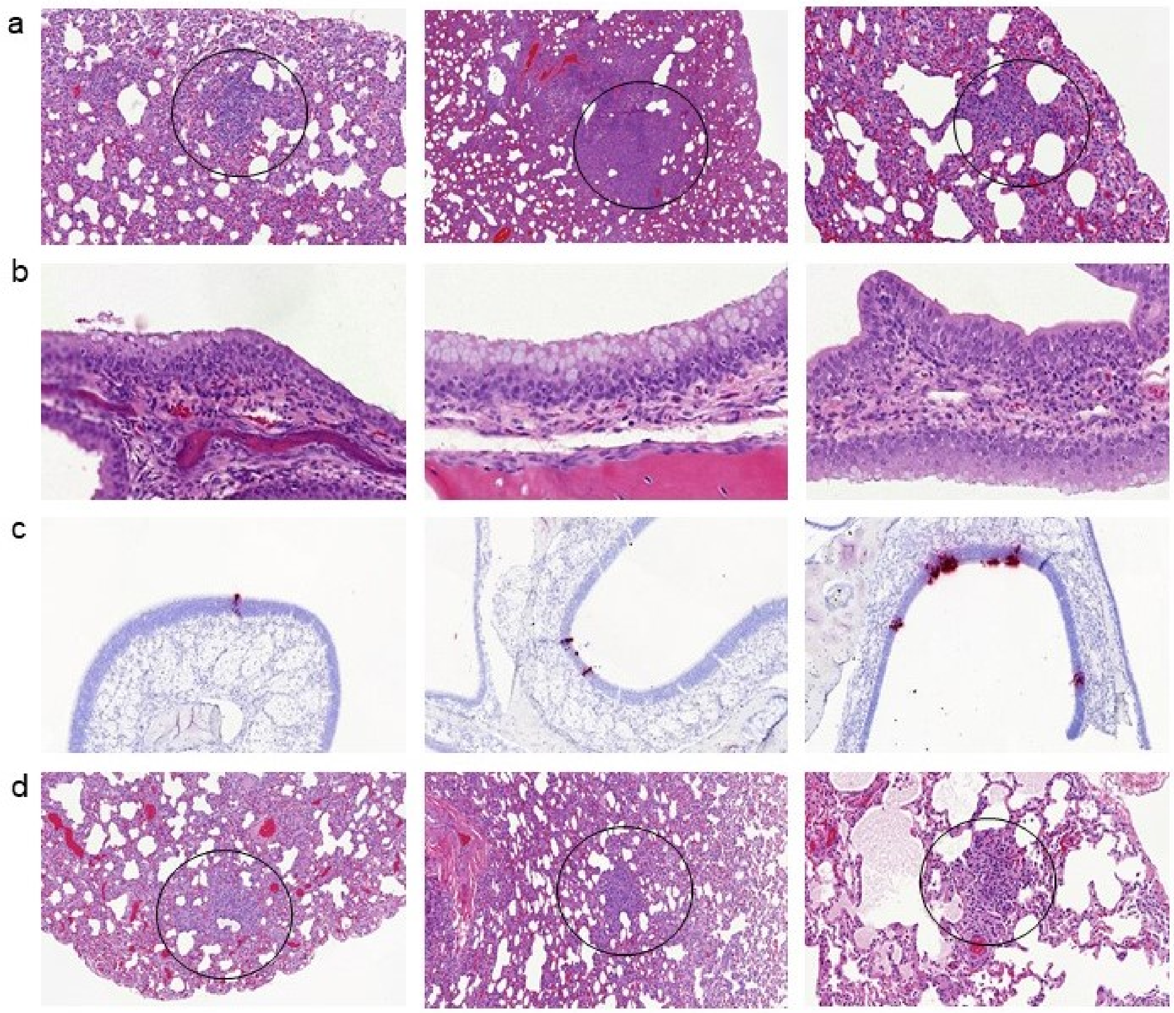


**Supplementary Figure 2. Lung and nasal cavity pathology.** Tissue samples from days 3 and 12-14 pc were fixed in 10% neutral buffered-formalin, processed to paraffin wax, sections cut at 4 µm thickness and stained with H&E; *in-situ* hybridisation (RNAscope) techniques were used to detect the presence of viral RNA in select nasal cavity samples. Overall, a number of microscopic changes attributable to SARS-CoV-2 virus were present at low severity in the lung and nasal cavity of ferrets in all groups at both time points. These comprised inflammatory changes involving infiltration of the alveoli, blood vessels (perivascular cuffing), and the airways of the lung, and the epithelia and underlying propria mucosa of the nasal cavity. At both time points, these changes were similar in appearance and severity in all groups. Overall, prominent differences in the severity of changes between animals in the INNA-051-treated test groups and the control group were not observed at either time point. At day 3 pc, occasional foci of cells staining positive for SARS-CoV-2 RNA were detected in the nasal cavity of animals in all group (tissues were not stained at the later time point). No difference in incidence of viral RNA positive cells in the nares was noted between groups.

A minimal selection of pathology is shown. Each panel of images shows from left to right an example tissue from the following groups; INNA-051 high dose group 1, INNA-051 low dose group 2, PBS control group 4.

a) Lung from day 3 pc (2/6 ferrets groups 1, 2 & 4); low-grade, foci of primarily mononuclear inflammatory cells, with occasional polymorph neutrophils, infiltrating alveolar walls and spaces (circles). An increase the severity of infiltration was seen in 1 of the 2 ferrets in group 2. HE. 330x, 140x and 500x magnification respectively.
b) Varying regions of the nares from day 3 pc (2/6 ferrets groups 1, 2 & 4); low-grade, neutrophilic cell infiltration in the epithelium and propria mucosa. HE. 820x, 820x and 600x magnification respectively.
c) Nasal cavity from day 3 pc (2/6 ferrets groups 1, 2 & 4;) rare, small foci of epithelial cells staining positive for SARS-CoV-2. RNAscope. All 400x magnification.
d) Lung from day 12-14 pc (4/6 ferrets groups 1, 2 & 4, 6/6 ferrets group 3 [not shown]); low-grade, foci of primarily mononuclear inflammatory cells, with occasional polymorph neutrophils, infiltrating alveolar walls and spaces (circles). HE. 270x, 263x and 410x magnification respectively.
